# Trypstatin as a Novel TMPRSS2 Inhibitor with Broad-Spectrum Efficacy Against Corona and Influenza Viruses

**DOI:** 10.1101/2025.01.14.632953

**Authors:** Jan Lawrenz, Lukas Wettstein, Armando Rodríguez Alfonso, Rayhane Nchioua, Pascal von Maltitz, Dan P.J. Albers, Fabian Zech, Julie Vandeput, Lieve Naesens, Giorgio Fois, Nico Preising, Emilia Schmierer, Yasser Almeida-Hernandez, Moritz Petersen, Ludger Ständker, Sebastian Wiese, Peter Braubach, Manfred Frick, Eberhard Barth, Daniel Sauter, Frank Kirchhoff, Elsa Sanchez-Garcia, Annelies Stevaert, Jan Münch

**Author notes:** Corresponding author: Jan Münch, Institute of Molecular Virology, Meyerhofstrasse 1, 89081 Ulm, Germany.

## Abstract

Respiratory viruses, such as SARS-CoV-2 and influenza, exploit host proteases like TMPRSS2 for entry, making TMPRSS2 a prime antiviral target. Here, we report the identification and characterization of Trypstatin, a 61-amino acid Kunitz-type protease inhibitor derived from human hemofiltrate. Trypstatin inhibits TMPRSS2 and related proteases, with IC_50_ values in the nanomolar range, comparable to the small molecule inhibitor camostat mesylate. *In vitro* assays demonstrated that Trypstatin effectively blocks spike-driven entry of SARS-CoV-2, SARS-CoV-1, MERS-CoV, and hCoV-NL63, as well as hemagglutinin-mediated entry of influenza A and B viruses. In primary human airway epithelial cultures, Trypstatin significantly reduced SARS-CoV-2 replication and retained activity in the presence of airway mucus. *In vivo*, intranasal administration of Trypstatin to SARS-CoV-2-infected Syrian hamsters reduced viral titers and alleviated clinical symptoms. These findings highlight Trypstatin’s potential as a broad-spectrum antiviral agent against TMPRSS2-dependent respiratory viruses.

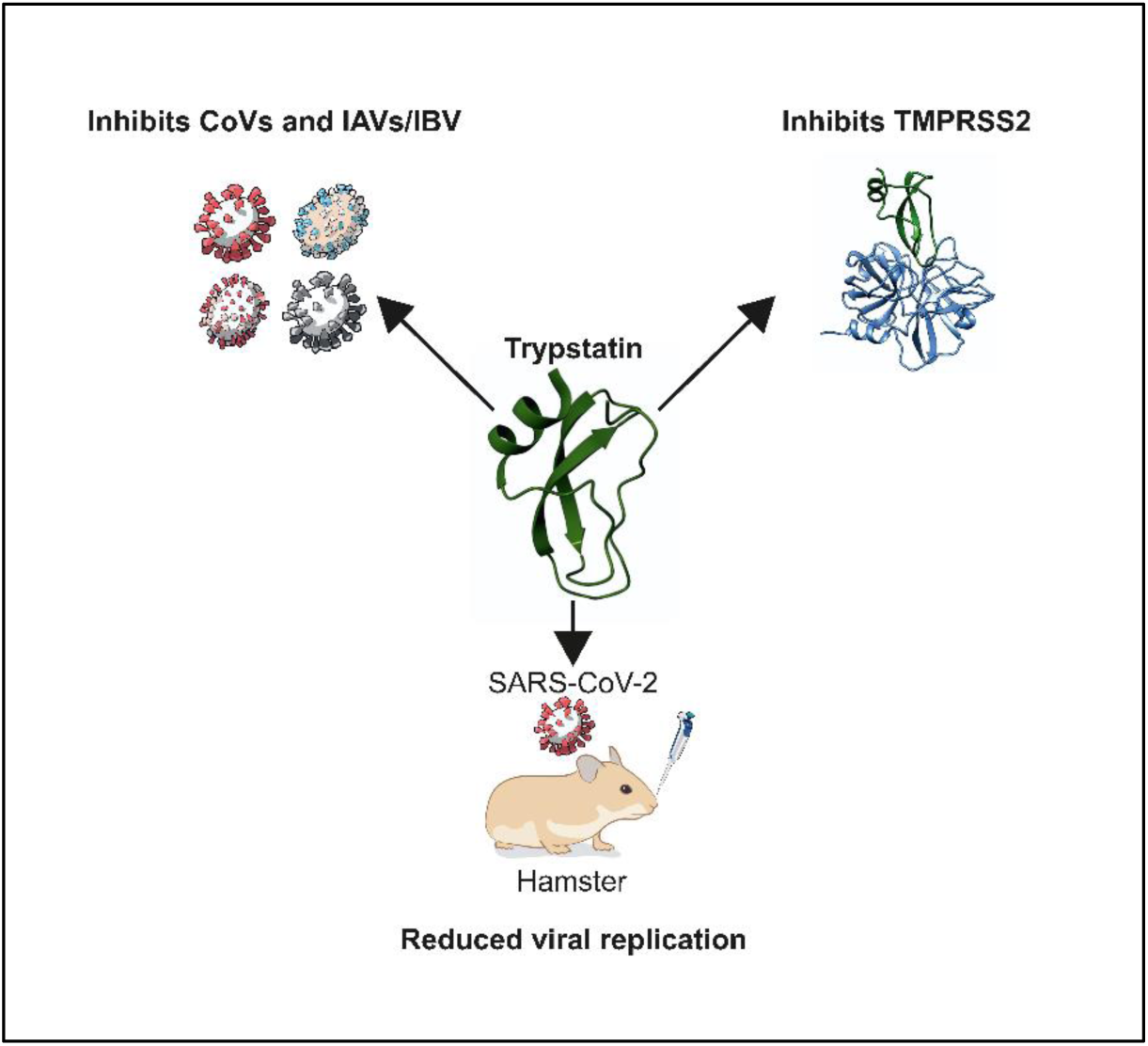

## 1. Introduction

Respiratory viruses, including influenza viruses and coronaviruses, pose a significant global health threat, causing substantial morbidity and mortality (1–4). These viruses are primarily transmitted through respiratory droplets, spreading rapidly within human populations. The COVID-19 pandemic caused by SARS-CoV-2 highlights the devastating potential of respiratory viruses to overwhelm healthcare systems and disrupt societies worldwide (5). Other respiratory viruses, such as influenza A and B, respiratory syncytial virus (RSV), and common cold-causing coronaviruses, contribute to annual respiratory disease burdens, with severe risks for vulnerable populations, including the elderly, young children, and immunocompromised individuals (6–8). Repeated zoonotic transmission of highly pathogenic respiratory viruses, such as SARS-CoV-1 (2002), MERS-CoV (2012), and SARS-CoV-2 (2019), underscore the urgent need for effective and broadly active antiviral therapies (9–11). Current virus-specific strategies targeting individual viral proteins or pathways are often limited in scope and require extensive development time for newly emerging viral pathogens (12). Broad-spectrum antivirals, which target shared components (13) or host factors critical for viral replication, represent a promising alternative. For respiratory viruses, host-directed approaches focusing on essential cellular enzymes, such as proteases, may inhibit viral entry and replication without exerting high selective pressure for resistance, making them attractive candidates for broad-spectrum therapy (14–16).

Many respiratory viruses utilize host cell proteases to initiate infection (17). One of the most studied proteases in this regard is the transmembrane protease serine 2 (TMPRSS2), which plays a critical role in viral entry by cleaving and activating viral envelope glycoproteins. For coronaviruses such as SARS-CoV-1, MERS-CoV and SARS-CoV-2, TMPRSS2 is essential for priming the spike (S) protein, facilitating fusion of the viral and host cell membranes and enabling viral entry (18–20). Similar mechanisms are exploited by influenza viruses, where proteolytic activation of the hemagglutinin (HA) protein is required for the virus to infect host cells (21–23). In addition to TMPRSS2, other proteases such as furin, trypsin-like serine proteases, and cathepsins can also be involved in viral entry processes, depending on the virus and cell type (17, 18, 20, 24–27). However, TMPRSS2 has gained particular attention for its role across multiple virus families (19, 21), making it a prime target for antiviral development.

To identify endogenous inhibitors of viral entry, we have utilized peptide-protein libraries derived from human body fluids and tissues as valuable sources of novel antivirals with unique modes of action (28–31). These libraries provide insights into the body’s natural defense mechanisms and offer potential therapeutic candidates. For example, VIRIP, a peptide isolated from human blood, blocks HIV-1 entry and an optimized derivative demonstrated significant efficacy in monotherapy trials by reducing viral loads in infected individuals (29, 32). Similarly, antitrypsin, identified in bronchoalveolar lavage (BAL) fluid, inhibits TMPRSS2 and SARS-CoV-2 entry, highlighting its potential as a therapeutic for both antitrypsin deficiency and respiratory viral infections (30). Here, we report the discovery of Trypstatin, a 61-mer Bikunin fragment and Kunitz-type protease inhibitor identified from a hemofiltrate-derived peptide-protein library. Trypstatin effectively inhibits TMPRSS2 and related proteases essential for the entry of SARS-CoV-2 and other respiratory viruses, including influenza A and B. Trypstatin demonstrates broad-spectrum antiviral activity, remains active in human airway mucus, reduces viral replication in primary airway cultures, and significantly lowers lung viral titers while improving clinical symptoms in SARS-CoV-2-infected Syrian gold hamsters. These properties highlight Trypstatin’s therapeutic promise as a host-directed antiviral agent against respiratory viruses.

## 2. Methods

### Cell culture

Caco-2 cells (human epithelial colorectal adenocarcinoma, kindly provided by Prof. Barth, Ulm University) were grown in DMEM supplemented with 100 U/ml penicillin, 100 µg/ml streptomycin, 1 mM sodium pyruvate, 2 mM L-glutamine, 1x non-essential amino acids and 20 % fetal calf serum (FCS). HEK293T-cells (ATCC® CRL-3216™) were cultivated in DMEM supplemented with 100 U/ml penicillin, 100 µg/ml streptomycin, 2 mM L-glutamin and 10 % FCS. Calu-3 cells (ATCC® HTB-55™) were cultured in Minimum Essential Medium Eagle (MEM) supplemented with 10 % FCS, 100 units/ml penicillin, 100 μg/ml streptomycin, 1 mM sodium pyruvate and 1x non-essential amino acids. LLC-MK-2 cells (kindly provided by Lia van der Hoek, Amsterdam, Netherlands) were cultured in Minimum Essential Medium Eagle (MEM) supplemented with 8 % FCS, 100 units/ml penicillin, 100 μg/ml streptomycin, 2 mM L-glutamine and 1x non-essential amino acids. Vero E6 cells (ATCC® CRL-1586™) were grown in DMEM supplemented with 100 units/ml penicillin, 100 μg/ml streptomycin, 2 mM L-glutamine, 1 mM sodium pyruvate, 1x non-essential amino acids, and 10% FCS. All cells were cultivated at 37°C in a 5 % CO_2_ humidified incubator. Cells were tested for mycoplasma contamination on a regular basis.

### Generation of human airway epithelial cells

Differentiated air-liquid interface cultures of human airway epithelial cells (HAECs) were generated from primary human basal cells isolated from airway epithelia as described previously (30). In brief, cells were expanded in Airway Epithelial Cell Basal Medium supplemented with Airway Epithelial Cell Growth medium SupplementPack (PromoCell). Medium was replaced every 2 days until 90 % confluency was reached and the HAECs were detached using DetachKIT (PromoCell) and seeded into 6.5 mm Transwell filters (Corning Costar). Filters were pre-coated with Collagen Solution (StemCell Technologies) overnight and irradiated with UV light for 30 min before 35,000 cells were seeded onto the apical side of each filter in 200 µl of growth medium supported by 600 µl of growth medium in the basolateral part. After 72-96 hours, when the cells reached confluence, apical medium was removed and basal medium was replaced by differentiation medium consisting of a 1:1 mixture of DMEM-H and LHC Basal (Thermo Fisher) supplemented with Airway Epithelial Growth Medium SupplementPack. Medium was replaced every 2 days. Removal of apical medium (air lifting) defined day 0 of air-liquid interface (ALI) culture. Cells were grown under ALI conditions until experiments were performed at day 25-27.

### Generation of the peptide/protein library from human hemofiltrate

The hemofiltrate library was generated in a previous work described by Lietz *et al.*(33). After the biological evaluation of the hemofiltrate library, the selected active fraction was subjected to several rounds of bioassay-guided purification, as described below, for the isolation of the biologically active compound.

### Subpurification of biologically active peptide library fractions

The dry material corresponding to the active fraction was dissolved in 10 ml of 95% A/5% B and fractionated by reversed-phase HPLC on a Luna C18 column (Phenomenex, USA) of dimensions 21.2 x 250 mm and particle size of 5 µm. The separation was performed at a flow rate of 12.23 ml/min using the gradient program (min/%B) 0/5, 5.09/5, 28.71/25, 49.55/50, and 70.39/100, being A, 0.1% TFA in water, and B, 0.1% TFA in acetonitrile. Elution was monitored online at a scan within the 190-600 nm range. Fractions were collected every minute and dried in a SpeedVac vacuum concentrator.

Second chromatographic step: the sample was dissolved in 1ml 95% A and 5% B and fractionated by reversed-phase HPLC on a Luna C18(2) column (Phenomenex, USA) of dimensions 10 x 250 mm and particle size of 5 µm. The separations were performed at a flow rate of 3.6 mL/min using the gradient program (min/%B) 0/5, 6/10, 60/50, and 65/100, being A, 0.1%TFA in water, and B, 0.1% TFA in acetonitrile. Elution was monitored online at 214, 220, 254, and 280 nm.

Third chromatographic step: the sample was dissolved in 1 ml 95% A/5% B and fractionated by reversed-phase HPLC on an Aeris Peptide XB-C18 column (Phenomenex, USA) of dimensions 4.6 x 250 mm and particle size of 3.6 µm. The separations were performed at a flow rate of 0.8 ml/min using the gradient program (min/%B) 0/3, 17/15, 55/75, and 65/100, being A, 0.1%TFA in water, and B, 0.1% TFA in acetonitrile. Elution was monitored online at 214, 220, 254, and 280 nm.

Fourth chromatographic step: The active fraction from the previous purification step was dissolved in 1.5 ml buffer A (see below) and subjected to cation-exchange chromatography on mono S HR 5/5 (Cytiva, USA) of dimensions 5 x 50 mm and particle size of 10 µm. The separation was performed at a flow rate of 1.5 ml/min using the gradient program (min/%B) 0/0, 5/0, 55/70, being A, ammonium formate with formic acid (0.016 M NH_4_^+^ + 0.024 M HCOO_-_/HCOOH) + 20% acetonitrile, pH 4.23, and B, ammonium formate with formic acid (0.8 M NH_4_^+^ + 1.22 M HCOO^-^/HCOOH) + 20% acetonitrile, pH 4.27. Elution was monitored online at 280 nm.

### Enzymatic digestion and LC-MS/MS sequencing

After purification, the sample containing the biologically active compound was treated with 5 mM DTT for 20 min at RT, carbamidomethylated with 50 mM iodoacetamide for 20 min at 37°C, and then quenched with 10 mM DTT. The carbamidomethylated sample was digested with trypsin (ThermoFisher #90059) at a 1:50 ratio (enzyme:protein) for 16 h at 37°C. A 15 µL-aliquot was analyzed by a nanoLC-Orbitrap Elite Hybrid mass spectrometry system (Thermo Fisher Scientific, Bremen, Germany) as previously described (34). Database searches were performed using PEAKs XPro (PEAKs studio 10.6) (35). For peptide identification, MS/MS spectra were correlated with the UniProt human reference proteome set. Carbamidomethylation at cysteine was set as a fixed modification while oxidation (M) and deamidation (NQ) were set as variable modifications. False discovery rates were set on the peptide level to 1%.

### Peptide synthesis and refolding

Trypstatin (AMBP 284-344, P02760), domain 2 (AMBP 287-337, P02760), and domain 1 (AMBP 231-281, P02760) aa sequences were retrieved from Uniprot (https://www.uniprot.org/). The peptides were synthesized by Synpeptide Co. Ltd., Shanghai, China. The three peptides were refolded to form their disulfide bridges. In brief, 10 mg of peptide was dissolved in 4 ml of denaturing buffer (50 mM Tris pH 8.5, 157 mM NaCl, 6 M GuHCl) containing 48 µl of a 1 M DTT solution before the mixture was incubated for 40 minutes at 40°C (600 rpm). Afterwards, 4 ml of denaturing buffer containing 59.8 mg glutathione disulfide (GSSG) was added and the mixture was dialyzed against 500 ml of renaturing buffer (50 mM Tris pH 8.5, 157 mM NaCl) for 16 hours at room temperature with magnetic stirring. The refolded sample was subjected to reversed-phase HPLC on an Aeris PEPTIDE XB-C18 column (Phenomenex, USA) of dimensions 10 x 250 mm and particle size of 5 µm, heated at 37°C. The separation was performed at a flow rate of 3 ml/min using the gradient program (min/%B) 0/5, 5/15, 65/45, 75/80, being A, 0.1%TFA in water, and B, 0.1% TFA in acetonitrile. Elution was monitored online at 280 nm. An Agilent 1100 Series (Agilent, USA) chromatographic system was used for the separation. Fractions were collected every minute and dried in a vacuum concentrator system (Labconco, USA). The anti-trypsin activity was found in the dominant peak of the chromatographic profile.

### Matrix-assisted laser-desorption ionization time-of-flight mass spectrometry (MALDI-TOF-MS) analysis of the refolded peptides

Refolded peptides were analyzed by an Axima Confidence MALDI-TOF MS (Shimadzu, Japan) in positive linear mode on a 384-spot stainless-steel sample plate. Spots were coated with 1 µl 5 mg/ml α-cyano-4-hydroxycinnamic acid (CHCA) previously dissolved in matrix diluent (Shimadzu, Japan), and the solvent was allowed to air dry. Then, a 0.5 µl sample or standard was applied onto the dry pre-coated well and immediately mixed with 0.5 µl matrix; the solvent was allowed to air dry. All spectra were acquired in the positive ion linear mode using a 337-nm N2 laser. Ions were accelerated from the source at 20 kV. One hundred profiles were acquired per sample, and 20 shots were accumulated per profile. The equipment was calibrated with a standard mixture in the TOFMixTM MALDI kit (Shimadzu, Japan). Measurements and MS data processing were controlled by the MALDI-MS Application Shimadzu Biotech Launchpad 2.9.8.1 (Shimadzu, Japan). High-purity fractions having the expected average mass (assuming the formation of three disulfide bridges) were selected for biological evaluation.

### Generation of lentiviral pseudoparticles

For generation of lentiviral pseudoparticles carrying coronavirus spike proteins 900,000 HEK293T-cells were seeded in 2 ml medium. On the next day, cells were transfected with 0.49 µg of pCMVdR8_91 (encoding a replication-deficient lentivirus), 0.49 µg pSEW-Luc2 (encoding a luciferase reporter gene, both kindly provided by Prof. Christian Buchholz, Paul-Ehrlich-Institute, Germany) and 0.02 µg of the respective glycoprotein encoding plasmid pCG1_SARS-2-SΔ18 (SARS-CoV-2 Hu-1, kindly provided by Prof. Stefan Pöhlman, DPZ Göttingen), pCG1_MERS-CoV and pCG1_SARS-CoV-1 (both kindly provided by Prof. Michael Schindler, Tübingen University) by mixing the plasmid DNA with TransIT®-LT1 Transfection Reagent at a 1:3 ratio in serum-free medium. After 20 min incubation at RT, the transfection mix was added dropwise to the cells. At 48 h post-transfection, pseudoparticle containing supernatants were harvested and clarified by centrifugation for 5 min at 1500 rpm. Virus stocks were aliquoted and stored at−80°C until use.

### Pseudoparticle inhibition assay

One day before transduction, 10.000 Caco-2 cells were seeded in DMEM supplemented with 10 % fetal calf serum, 2 mM L-glutamine, 100 U/I penicillin, 100 µg/ml streptomycin, 1x non-essential amino acids and 1 mM sodium pyruvate in a 96-well flat bottom plate. The next day medium was replaced by 60 μl serum-free growth medium and cells were treated with serially diluted compounds for 30 min at 37°C followed by transduction of cells with 20 μl of respective lentiviral pseudoparticles. Transduction rates were assessed by measuring luciferase activity in cell lysates at 48 h post-transduction with a commercially available kit (Luciferase Assay System, Promega) in an Orion II microplate reader with simplicity 4.2 software. Values for H_2_O controls were set to 100 % pseudoparticle entry.

### Screening hemofiltrate library for inhibitors of SARS-CoV-2 spike-driven entry

10,000 Caco-2 cells were seeded in 100 µl respective medium in a 96 well flat-bottom plate. The following day, medium was replaced by 60 µl of serum-free medium before 10 µl of the solubilized peptide containing fraction (in dH_2_O) was added and cells were transduced with 20 µl of luciferase-encoding lentiviral pseudoparticles carrying the SARS-CoV-2 Hu-1 spike. Transduction rates were assessed 48h post transduction by measuring luciferase activity in cell lysates as already described above.

### Viral strains and propagation

HCoV-NL63 was obtained from Lia van der Hoek, Amsterdam University and propagated in LLC-MK2 cells as described before (36, 37). SARS-CoV-2 B.1 D614G was retrieved from European Virus Archive (BetaCoV/Netherlands/01/NL/2020 #010V-03903). SARS-CoV-2 BA.1 (Omicron variant, lineage B.1.1.529, GSAID reference hCoV-19/Netherlands/NH-RIVM-71084/2021) was retrieved from European Virus Archive, Omicron BA.5 (B.1.1.529) was kindly provided by Prof. Dr. Florian Schmidt (University of Bonn), and propagated as described previously (30). Influenza A/Virginia/ATCC3/2009 (Virg09; A/H1N1 subtype) was purchased from ATCC® (VR-1737™), Influenza A/Victoria/361/11 virus (Vict11; A/H3N2) was provided by G. Rimmelzwaan (Rotterdam, The Netherlands) and Influenza B/Ned/537/05 (Ned05; B/Yamagata lineage) was generously donated by R. Fouchier (Rotterdam, The Netherlands). Influenza Viruses were expanded in 10-day-old embryonated chicken eggs. For virus titration, a virus dilution series was added in quadruple to Calu-3 cells. At day 3 post infection, virus positivity was assessed by immunostaining for viral nucleoprotein NP as described in the antiviral assay section. Virus titers for all viruses were expressed as the 50% cell culture infective dose (TCID_50_), calculated according to the method of Reed and Muench (38).

### SARS-CoV-2 inhibition assay in Caco-2 and Calu-3 cells

To assess inhibitory activity of Trypstatin against SARS-CoV-2 infection, 25000 Caco-2 cells were seeded in a 96-well flat-bottom plate before on the following day the cells were treated for 30 minutes with serial dilutions of Trypstatin or Camostat mesylate. Following this, cells were infected with SARS-CoV-2 B.1 D614G (MOI 0.0005) or the Omicron BA.1 variant (MOI 0.23). After 24h, medium was removed and cells detached using Trypsin-EDTA before cells were fixed in a final concentration of 4 % paraformaldehyde for 30 minutes at room temperature. Cells were washed with PBS and stained for flow cytometry. Briefly, cells were incubated with anti-SARS-CoV-2 nucleocapsid antibody (SinoBiological #40143-MM05) diluted 1:1000 in Buffer B (Fix and Perm, MuBio Nordic) for 40 minutes at 4°C. Cells were washed twice with FACS buffer (1 % FCS in PBS) and incubated with Alexa Flour 488 labeled anti-mouse antibody (Thermo Fisher #A32723) diluted 1:400 in FACS buffer for 30 minutes at 4°C before the cells were washed again twice with FACS buffer. Analysis was performed using a CytoflexLX flow cytometer (Beckmann Coulter) with CytExpert 2.3 software.

To assess inhibitory activity of Trypstatin against SARS-CoV-2 replication, 140000 Calu-3 cells were seeded in a 24 well plate. The next day, medium was replaced and cells treated with Trypstatin or camostat mesylate for 30 min followed by infection with SARS-CoV-2 Omicron BA.5 (MOI 0.001). 3 hours post infection (hpi), cells were washed three times with PBS and growth medium supplemented with respective compound was added. Supernatants were samples after the washing step (day 0) and at 1, 2 and 3 days post-infection (dpi). At each sampling point, medium was removed completely and fresh medium supplemented with compounds was added. For analysis of SARS-CoV-2 replication, viral genome copies were quantified in supernatant samples. To this end, viral RNA was isolated using the QIAmp Viral RNA Mini Kit (Qiagen). A reaction mix consisting of 1x Fast Virus 1-Step Mastermix (Thermo Fisher #4444436), 0.5 µM of each Taqman primer targeting SARS-CoV-2-ORF1b-nsp14 (fwd primer: 5’TGGGGYTTTACRGGTAACCT-3’; rev primer: 5’-AACRCGCTTAACAAAGCACTC-3’), 0.25 µM probe (5’-FAM-TAGTTGTGATGCWATCATGACTAG-TAMRA-3’) and 5 µl of isolated viral RNA was prepared and applied to the following cycling conditions: 1 cycle reverse transcription (50°C, 300 s) and RT inactivation (95°C, 20 s), 40 cycles of denaturation (95°C, 5 s) and extension (60°C, 30 s) in a StepOnePlus qPCR cycler (Applied Biosystems) with StepOne Software 2.3. An in-house RNA standard based on a synthetic SARS-CoV-2 RNA standard (Twist Bioscience #102024) was used to determine genome copies from C_t_ values.

### SARS-CoV-2 infection of human airway epithelial cells (HAECs)

Right before infection, the apical surface of HAECs grown on Transwell filters at the air-liquid interface were washed three times with pre-warmed PBS to remove mucus. Subsequently, 10 µM Trypstatin, 80 µM camostat mesylate or PBS were added onto the apical surface for 10 minutes before viral inoculum (SARS-CoV-2 Omicron BA.5, MOI 0.5) was added for two hours. Following this, the inoculum and compound was removed, cells washed with pre-warmed PBS and further cultured at the air-liquid interface. Accumulated mucus was washed off daily with pre-warmed PBS. Two days post-infection, mucus was removed and PBS added to the apical side for 30 minutes before the sample was subjected to TCID_50_ titration on Vero E6 cells. TCID_50_ was calculated 6 days post-infection according to Reed and Muench (38).

### SARS-CoV-2 infection of Syrian gold hamsters

Twelve 8-week-old male Syrian gold hamsters obtained from Janvier, France, were randomized into two groups (6+6). Group 1 received vehicle (PBS) and group 2 received 0.3 mg/kg Trypstatin. On day 1, animals were subjected to two intranasal administrations of Trypstatin or PBS, 6 hours apart. Three hours after the first administration, all animals were subjected to intranasal inoculation with 2×10^4^ PFU SARS-CoV-2 Delta strain. On day 2 and 3 animals received one intranasal administration of Trypstatin or PBS. All administrations were done under 3,5% isoflurane anaesthesia. 72 hours post-infection animals were euthanized and bronchoalveolar lavage was collected. Viral load and TCID_50_/ml of BAL was performed as described above. Clinical symptoms of animals were recorded daily according to a modified Irwin screen. Animal experiments were carried out by Scantox in Solna, Sweden under BSL3 conditions.

### HCoV-NL63 inhibition assay

25,000 Caco-2 cells were seeded in a 96-well flat-bottom plate before the following day, cells were treated for 30 minutes with serial dilutions of compounds in PBS and the cells were infected with a MOI of 0.018 and incubated at 33°C. 72 hpi, cells were detached and fixed in a final concentration of 4 % PFA for 30 minutes. Subsequently, cells were washed with PBS and stained for flow cytometry analysis with anti-NL63-nucleocapsid antibody (SinoBiological #40641-T62) diluted 1:5000 in Buffer B (Fix and Perm, MuBio Nordic) for 40 minutes at 4°C. Cells were washed twice with FACS buffer (1 % FCS in PBS) and incubated with Alexa Flour 647 labeled anti-rabbit antibody (Cell Signaling #4414) diluted 1:5000 in FACS buffer for 30 minutes at 4°C before the cells were washed again twice with FACS buffer. Analysis was performed using a CytoflexLX flow cytometer (Beckmann Coulter) with CytExpert 2.3 software.

### Influenza virus inhibition assay

Antiviral activity in IAV-or IBV-infected Calu-3 cells was determined as described previously (23), with minor modifications. Calu-3 cells were seeded in black 96-well plates at 40,000 cells per well. One day later, the cells were exposed to serial compound dilutions for 30 minutes. Next, the cells were infected with IAV or IBV at 100xTCID_50_ per well. At two hpi, the cells were washed to remove excess virus, after which fresh compound dilutions were added. After three days of incubation at 35 °C, the cells were immunostained for viral NP. First, the cells were fixed with 2% paraformaldehyde and permeabilized with 0.1% Triton X-100. Next, they were stained with anti-NP antibody (for IAV: 3IN5 InA108 [HyTest] at 1:2,000; for IBV: 3IF18 R2/3 [HyTest] at 1:2,000). Goat anti-mouse IgG Alexa Fluor 488 (A11001 at 1:500; Invitrogen) was used as secondary antibody and cell nuclei were stained with Hoechst (Thermo Fisher Scientific). Nine images per well were analyzed, using a CellInsight CX5 high-content imaging platform (Thermo Scientific) to determine the numbers of total (Hoechst) and infected (NP) cells.

### Molecular dynamics simulations

The coordinates of trypstatin were obtained from the bikunin structure (PDB ID: 1BIK (39), residues 80-113). For TMPRSS2, an AlphaFold (40, 41) model (UniProt ID: O15393) was used for the catalytic domain (residues 256-491). To generate the starting structure of the TMPRSS2/trypstatin complex, both molecules were superimposed onto the complex of bovine chymotrypsin with a Kunitz-type serine protease inhibitor (PDB ID: 3M7Q (42)). The system was prepared with CHARMM-GUI (43) to run classical molecular dynamics (MD) simulations. The systems were placed in a cubic simulation box with 10 Å of padding. The box was solvated with TIP3P water model (44) and NaCl at 150 mM. After regular minimization and equilibration, 5 replicas of production runs were performed during 500 ns each for TMPRSS2/trypstatin. The simulations were run with GROMACS (45) using the CHARMM36m force field (46). For the analysis of the simulations, the last 300 ns (1.5 μs in total) were employed. Clustering analysis with the VMD (47) clustering plugin (https://github.com/luisico/clustering) was performed on the RMSD of the backbone of the complex, with a cutoff = 3 Å. The centroid of the first cluster was taken as a representative structure. The PPI-Affinity tool (48) was employed to compute the binding affinity of the TMPRSS2/trypstatin complex, as well as the binding affinities of the TMPRSS/anti-thrombin (49) and TMPRSS2/anti-trypsin (30) complexes.

### Recombinant TMPRSS1, TMPRSS2 and TMPRSS14 activity assay

For assessing the inhibition of recombinant human TMPRSS’s, 25 µl of serially diluted compound was incubated with 25 µl of recombinant enzyme (TMPRSS2: 2 µg/ml, LSBio #LS-G57269; TMPRSS1: 0.02 µg/ml, R&D #11416; TMPRSS14: 0.2 µg/ml, #3946) in assay buffer (TMPRSS2: 50 mM Tris-HCl, 0.154 mM NaCl pH 8.0; TMPRSS1/14: 50 mM Tris, 0.05 % Brij35, pH 9) for 15 min at 37 °C. In a next step, 50 µl of 20 µM BOC-Gln-Ala-Arg-AMC protease substrate (Bachem #4017019) in the case of TMPRSS2 and TMPRSS14 or 50 µl of 150 µM BOC-Gln-Arg-Arg-AMC (PeptaNova) in the case of TMPRSS1 was added and incubated for 2 h for TMPRSS2, or 5 mins in the case of TMPRSS1/14 at 37°C. Fluorescence intensity was measured after 2 h at an excitation wavelength of 380 nm and emission wavelength of 460 nm in a SynergyTM H1 microplate reader (BioTek) with Gen5 3.04 software.

### Recombinant neutrophil elastase activity assay

Neutrophil Elastase activity was measured by mixing 25 µl of compound with 25 μl of 2 ng/μl recombinant neutrophil elastase (Merck Millipore #324681) in assay buffer (50 mM Tris, 1 M NaCl, 0.05 % (w/v) Brij-35, pH 7.5) for 15 min at 37°C. Next, 50 μl of 200 μM of MEOSUC-Ala-Ala-Pro-Val-AMC substrate (Bachem #4005227) was added and incubated at 37°C. Fluorescence intensity was measured after 5 minutes at an excitation wavelength of 380 nm and emission wavelength of 460 nm in a SynergyTM H1 microplate reader (BioTek) with Gen5 3.04 software.

### Recombinant kallikrein 5 activity assay

Kallikrein 5 activity was measured by mixing 25 µl of compound with 25 µl of 2µg/ml of recombinant enzyme (R&D #1108) in assay buffer (100 mM NaH2PO4, pH 8) for 15 min at 37°C before the addition of 50 µl of BOC-Val-Pro-Arg-AMC substrate (R&D #ES011). Mixture was incubated for 5 minutes at 37°C before fluorescence intensity was measured at an excitation wavelength of 380 nm and emission wavelength of 460 nm in a SynergyTM H1 microplate reader (BioTek) with Gen5 3.04 software.

### Cellular TMPRSS activity assay

20,000 HEK293T-cells were seeded in a 96-well flat bottom plate. The next day, cells were transfected with 100 ng of an expression plasmid (Backbone: Twist Bioscience, EF1alpha mammalian expression vector) encoding the respective transmembrane serine protease per well using LT1 transfection reagent (TMPRSS2 human NCBI RefSeq: NP_005647.3; TMPRSS2 syrian gold hamster NCBI RefSeq: XP_012971683.1; TMPRSS11D NCBI RefSeq: NP_004253.1; TMPRSS13 NCBI RefSeq: NP_001070731.1). In short, plasmid DNA was mixed in a 1:3 ration with LT1 in serum free medium and incubated for 20 minutes before the mixture was applied dropwise onto the cells. 16h post-transfection, medium was replaced by serum-free medium and cells were treated with serial dilution of compounds in PBS. After 15 minutes of incubation at 37°C, 100 µM of respective protease specific substrate (TMPRSS2: Boc-QAR-AMC, Bachem # 4017019; TMPRSS11D: Boc-FSR-AMC, Bachem # 4012340; TMPRSS13: Pyr-RTKR-AMC, Bachem #4018149) was added and incubated for 2 h at 37°C. Fluorescence intensity was recorded at an excitation wavelength of 380 nm and emission wavelength of 460 nm in a Synergy™ H1 microplate reader (BioTek) with Gen 5 3.04 software. Values were background corrected for non-transfected HEK293T-cells.

### Cathepsin activity assay

Cathepsin enzyme activity assays were performed as described before (49). In brief, serially diluted compounds were mixed with recombinant cathepsin L (R&D #952-Cy) in assay buffer (50 mM MES, 5 mM DTT, 1 mM EDTA, 0.005% Brij35, pH 6) or with cathepsin B from human placenta (Sigma Aldrich #C0150) in assay buffer (105.6 mM KH_2_PO_4_, 14.4 mM Na_2_HPO_4_, 1.2 mM EDTA, 0.07% Brij35, 2.4 mM L-cysteine, pH 6). The protease-inhibitor mixtures were incubated for 10 min at RT before addition of the respective fluorogenic reporter substrate Z-Leu-Arg-AMC (for cathepsin L; Bachem #4034611) or Z-Arg-Arg-AMC (for cathepsin B; Bachem #4004789). Final concentrations were as follows: Cathepsin L 0.01 μg/ml, cathepsin B 1 μg/ml, Z-Leu-Arg-AMC 40 μM or Z-Arg-Arg-AMC 30 μM. Fluorescence intensity was recorded at an excitation wavelength of 380 nm and emission wavelength of 460 nm after 25 min at 37°C (cathepsin L) or after 60 min at 40°C (cathepsin B) in a Synergy™ H1 microplate reader (BioTek) with Gen 5 3.04 software.

### Furin activity assay

Furin activity assay was conducted as described previously (49). In brief, Trypstatin was mixed with recombinant furin (R&D #1503-SE-010, final concentration 1.5 ng/µl) in assay buffer (25 mM Tris, 1 mM CaCl_2,_ pH 9) for 10 min at 37°C before the addition of 20 µM succinyl-modified QTNSPRRAR-AMC substrate. Fluorescence intensity was recorded in 2 min intervals for 5 h at an excitation wavelength of 355 nm and emission wavelength of 460 nm at 37°C in a Cytation 3 (BioTek) microplate reader. Area under the curve values were normalized to the furin control.

### Reagents

Camostat mesylate (#SML0057) and E-64d (#E8640) were obtained from Merck. Antithrombin (Anbinex®) and α1-antitrypsin (Prolastin®) were obtained from Grifols. Hexa-D-Arginine was purchased from Biozol (#MCE-HY-P1028), Aprotinin from Sigma Aldrich (#A1153), Nirmatrelvir from Selleckchem (#PF-07321332) and Baloxavir acid from MedChem Express (#HY-109025A).

### Cytotoxicity assay

To assess the cytotoxicity of the inhibitors used, 10,000 Caco2 cells were seeded in a 96-well flat bottom plate. The next day, the medium was replaced and cells were treated with serial dilutions of compounds or PBS as control. After 48 h, cell viability was assessed by measuring ATP levels in cell lysates with a commercially available kit (CellTiter-Glo®, Promega) in an Orion II microplate reader with Simplicity 4.2 software.

### Statistical Analysis

Unless stated otherwise, analysis was performed using GraphPad Prism version 10.3.1. Calculation of IC_50_ values via nonlinear regression was performed using normalized response-variable slope equation. For statistical analysis, two-tailed Mann-Whitney test was used as indicated in the respective figure legend. The number of animals or repeated experiments is stated in each figure legend.

## 3. Results

### Identification of Trypstatin as inhibitor of SARS-CoV-2

To identify endogenous factors that inhibit SARS-CoV-2 infection, we generated and screened a peptide-protein library derived from human hemofiltrate for inhibitors of spike-driven entry. Similar to previous approaches (30, 31), peptides and small proteins were extracted from about 1000 liter of hemofiltrate by chromatographic means into eight pH pools, each consisting of 55 fractions. These fractions were then tested in human epithelial colorectal carcinoma (Caco-2) cells using luciferase-encoding lentiviral pseudoparticles carrying the SARS-CoV-2 spike protein. Strong inhibition of viral entry was observed in pH pool 3, with fraction 19 showing the highest activity (Figure 1A).

**Figure 1:**
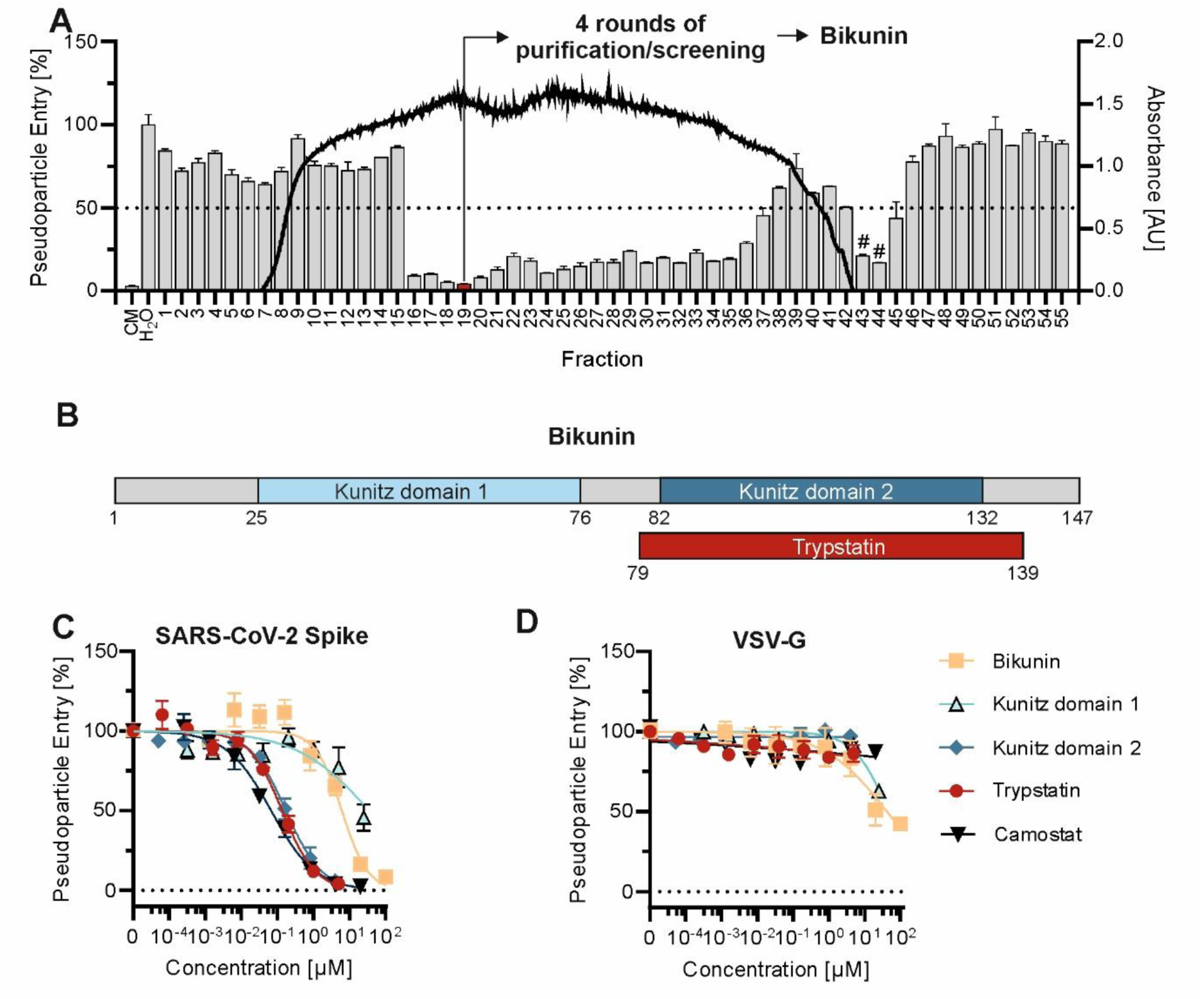
High-throughput screening for endogenous inhibitors of SARS-CoV-2 entry. **(A)** Caco-2 cells were pre-treated with peptide/protein containing fractions of a hemofiltrate library before transduction with luciferase-encoding lentiviral pseudoparticles harboring the SARS-CoV-2 Hu-1 spike. Transduction rates were assessed 48 h later by measuring luciferase activity in cell lysates. Columns represent pseudoparticle entry (expressed as % versus H2O-treated condition) and black line absorbance at 280 nm of the corresponding fraction. The serine protease inhibitor camostat mesylate (CM) was included as reference inhibitor. Hit fraction 19 was subpurified and screened again (Supplemental Figure 1). Shown are mean values of one experiment performed in triplicates ± SEM. CM = Camostat mesylate. # indicates cellular toxicity as observed by light microscopy **(B)** Schematic representation of serine protease inhibitor Bikunin with protease inhibitor domains from the kunitz-type. **(C-D)** Caco-2 cells were pre-treated with serial dilution of compounds before transduction with luciferase-encoding lentiviral pseudoparticles harboring the SARS-CoV-2 Hu-1 spike or VSV-G glycoprotein. Transduction rates were assessed 48 h later by measuring luciferase activity in cell lysates. Shown are mean values of three independent experiments performed in triplicates ± SEM.

Fraction 19 underwent several rounds of chromatographic purification and screening, ultimately yielding antivirally active fractions (30–32) after four purification cycles (Supplemental Figure 1). LC-MS/MS analysis of these active fractions identified a large fragment of the Kunitz-type serine protease inhibitor inter-alpha-trypsin inhibitor light chain, known as Bikunin (Uniprot P02760, AMBP_HUMAN), spanning residues 221-349. Bikunin contains two protease inhibitor domains: Bovine Pancreatic Trypsin Inhibitor (BPTI)/Kunitz-type domain 1 (residues 231-281) and domain 2 (residues 287-337) (Figure 1B). Previously, Kido *et al*. reported the isolation of a Bikunin fragment from rat mast cells, named Trypstatin, which is slightly larger than the second Kunitz domain (50, 51). Based on this discovery, a human homologue was inferred through sequence homology, corresponding to residues 284-344 of Bikunin, and also named Trypstatin.

To determine the inhibitory potential of full-length Bikunin or of its individudal Kunitz domains, we tested commercially available Bikunin (purified from urine), along with chemically synthesized and refolded Kunitz domains 1, 2, and the Trypstatin fragment (MS-spectra in Supplemental Figure 2). To this end, Caco-2 cells were treated with serial dilutions of the compounds before transduction with SARS-CoV-2 spike pseudoparticles, or VSV-G pseudoparticles as controls. Full-length Bikunin partially suppressed SARS-CoV-2 entry (IC_50_ 6.1 µM) but also showed non-specific effects against VSV-G-mediated entry (Figure 1C-D). By contrast, Kunitz domain 2 and the related Trypstatin fragment specifically blocked SARS-CoV-2 entry in the nanomolar range with IC_50_’s of 170 and 136 nM, respectively, without affecting VSV-G-driven entry, similar to the small molecule inhibitor camostat mesylate (CM). No cytotoxicity was observed for any compound at the tested concentrations (Supplemental Figure 3). Although domain 2 is slightly shorter and almost equally active, we focused our further experiments on the Trypstatin fragment due to its presumed occurrence in the human body.

### Trypstatin inhibits TMPRSS2 and related transmembrane serine proteases

Given that SARS-CoV-2 entry relies on proteolytic cleavage of the spike protein, we hypothesized that Trypstatin, as a Kunitz-type protease inhibitor, might target key spike-processing proteases, specifically TMPRSS2, Furin, and/or Cathepsins L and B (24, 52, 53). To test this hypothesis, we conducted protease activity assays with recombinant and cellular proteases using fluorogenic substrates. Our results showed that Trypstatin is a potent inhibitor of TMPRSS2 with IC_50_ values of 3.9 nM (recombinant TMPRSS2, Figure 2A) and 7.9 nM (cellular TMPRSS2, Figure 2B). Trypstatin did not inhibit Cathepsins L and B (Figure 2C, D) or Furin (Supplemental Figure 4). Interestingly, Trypstatin also effectively inhibited related transmembrane serine proteases TMPRSS11D (IC_50_: 24.8 nM, Figure 2E) and TMPRSS13 (IC_50_: 17.6 nM, Figure 2F), which are known to cleave the SARS-CoV-2 spike protein and glycoproteins of other respiratory viruses (14, 21, 26). Notably, Trypstatin’s potency against TMPRSS2, TMPRSS11D, and TMPRSS13 was similar or superior to that of the small molecule inhibitor camostat mesylate, positioning Trypstatin as the most potent naturally occurring TMPRSS2 inhibitor identified to date. To further elucidate the anti-proteolytic profile of Trypstatin, we extended our investigation to proteases beyond the well-established glycoprotein priming proteases. Specifically, we examined its inhibition of two additional transmembrane serine proteases, TMPRSS1 (Hepsin) and TMPRSS14 (Matriptase), as well as the serine proteases elastase and kallikrein, which are associated with inflammation (54–56). Trypstatin inhibited TMPRSS1, neutrophil elastase and kallikrein 5 with activities in the nanomolar range, while having no effect against TMPRSS14 (Figure 2G-J). In summary, Trypstatin is a potent inhibitor of TMRPSS enzymes involved in viral entry, being particularly potent against TMPRSS2 but also additionally blocks proteases involved in pro-inflammatory cascades, albeit with higher IC_50_’s (Figure 2K).

**Figure 2:**
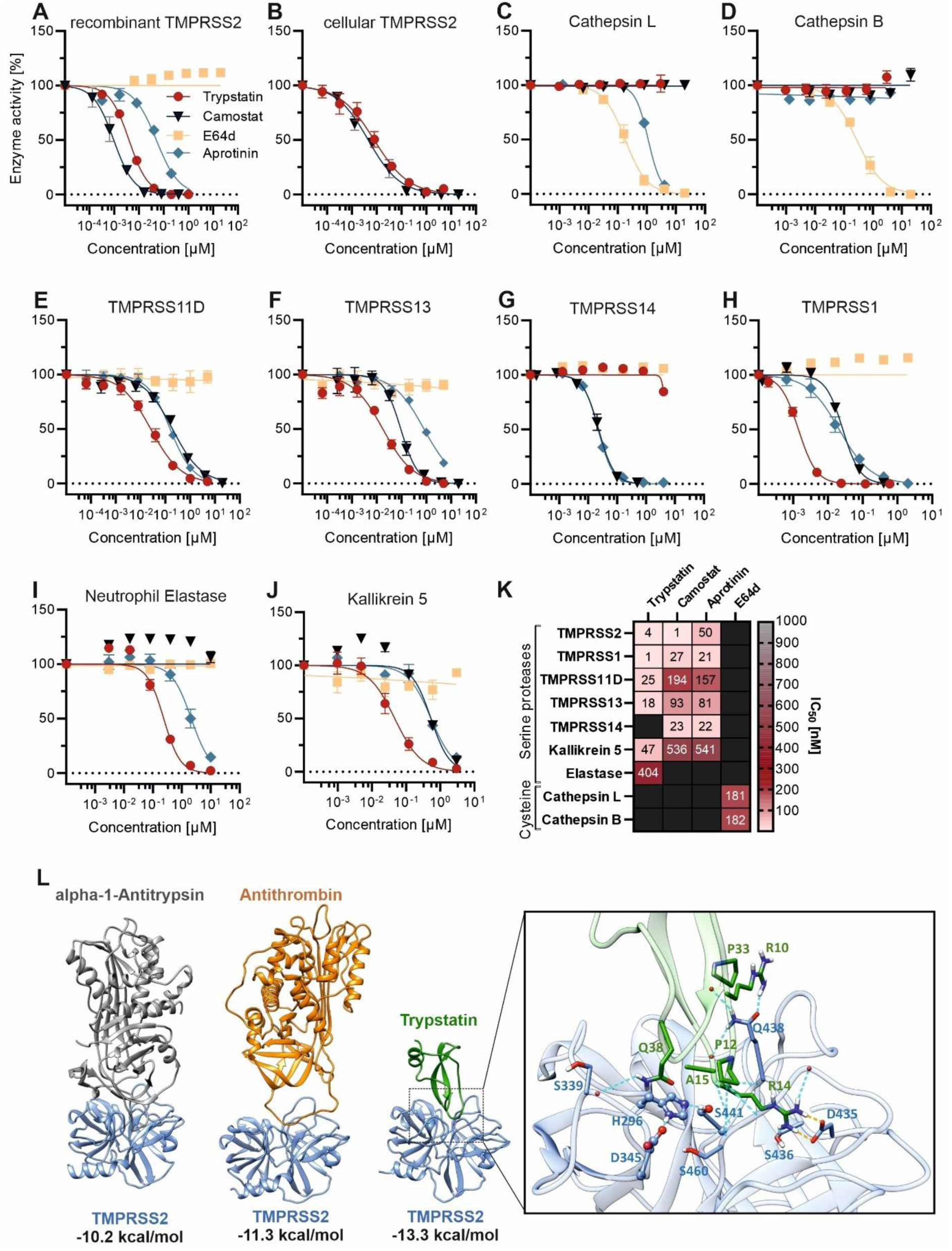
Trypstatin inhibits TMPRSS2 and related proteases. (A, C-D, G-J) Serial dilutions of Trypstatin, serine protease inhibitor camostat mesylate, kunitz inhibitor aprotinin and cysteine protease inhibitor E64-d were mixed with the respective enzyme before addition of a fluorogenic reporter substrate. Fluorescence intensity was measured at an excitation wavelength of 380 nm and emission wavelength of 460 nm. **(B, E-F)** HEK293T-cells were transfected with TMPRSS2-, TMPRSS11D- and TMPRSS13-encoding plasmid or mock control, respectively, before treatment with serial dilutions of inhibitors and the addition of a fluorogenic reporter substrate. Values were corrected for the signal of mock-transfected HEK293T-cells. Shown are mean values of three independent experiments performed in triplicates ± SEM. **(K)** Heat-map summarizing the IC50 values of A-J. **(L)** Binding affinity prediction between protease inhibitors and TMPRSS2 with PPI-Affinity. The complexes are representative structures obtained by molecular dynamics simulations. Zoom: S1 pocket of TMPRSS2. Hydrogen bonds are shown in light blue, salt bridges in orange. The catalytic residues (H296, D435 and S441) are depicted in Corey-Pauling-Koltun representation.

Molecular dynamics simulations revealed that the binding of Trypstatin to TMPRSS2 is primarily mediated through molecular interactions at the S1 pocket of TMPRSS2, a specificity site within the protease that binds critical residues of its substrates (Figure 2L). The analysis of the last 300 ns of all the replicas (1.5 µs in total) showed that Trypstatin’s P1-positioned arginine (R14), a residue specifically aligned to interact with the S1 pocket, forms a stable salt bridge with a conserved aspartic acid residue (D435) in TMPRSS2. This binding configuration blocks substrate access to the protease active site, thereby inhibiting enzymatic function. In protease-substrate interactions, the S1 pocket is a well-defined area on the protease that accommodates the P1 residue of the substrate (or inhibitor), with the S1-P1 pairing often dictating specificity and binding strength (57). Here, the P1 arginine of Trypstatin is critical for binding affinity, as its positive charge allows it to engage on electrostatic interactions with D435 and hydrogen bond interactions with S436 at the S1 pocket of TMPRSS2 (Figure 2L). Furthermore, binding affinity estimations (48) using representative structures from the MD simulations of trypstatin (see Supplemental Figure 5 for more details), anti-thrombin (49), and alpha-1-antitrypsin (30) in complex with TMPRSS2 indicate that the affinity of Trypstatin for TMPRSS2 (binding energy ∼ -13.3 kcal/mol) surpasses that of other endogenous inhibitors alpha-1-antitrypsin and antithrombin, with binding affinity energies of -11.3 kcal/mol and - 10.2 kcal/mol, respectively (Figure 2L).

### Trypstatin inhibits SARS-CoV-2 *in vitro* and *in vivo*

After establishing the mechanism of action of Trypstatin, we evaluated its inhibitory potential against SARS-CoV-2 spike-driven entry relative to known protease inhibitors reported to block TMPRSS2 and SARS-CoV-2 infection. As shown in Figure 3A, Trypstatin demonstrated lower IC_50_ values (111.9 nM) than α-1-antitrypsin (IC_50_ ∼27.6 µM), antithrombin (IC_50_ ∼1.5 µM), and aprotinin (IC_50_ 420.4 nM), achieving inhibition similar to the small-molecule inhibitor Camostat mesylate (IC_50_ 76.8 nM). To further confirm its potency, we conducted inhibition assays in Caco-2 cells infected with authentic SARS-CoV-2 B.1 D614G. Trypstatin showed an IC_50_ of 26 nM (Figure 3B, left). The Omicron BA.1 variant was also effectively inhibited, albeit at a slightly higher IC_50_ of 124 nM (Figure 3B, right). Treatment with 10 µM Trypstatin reduced replication of SARS-CoV-2 BA.5 in human lung cells (Calu-3) over the course of 72 hours by up to four orders of magnitude (Figure 3C). Analysis of cumulative viral RNA copies showed a more than 1000-fold (99.9%) reduction with Trypstatin treatment (Figure 3D). Next, we evaluated Trypstatin’s efficacy in a more physiologically relevant model using fully differentiated primary human airway epithelial cells (HAECs) grown at the air-liquid interface. HAECs from three donors were treated with 10 µM Trypstatin or 80 µM camostat mesylate while exposed to SARS-CoV-2 Omicron BA.5. A single apical application of Trypstatin resulted in a 68- to 684-fold reduction in viral loads in apical wash samples two days post-infection, depending on the donor (Figure 3E).

**Figure 3:**
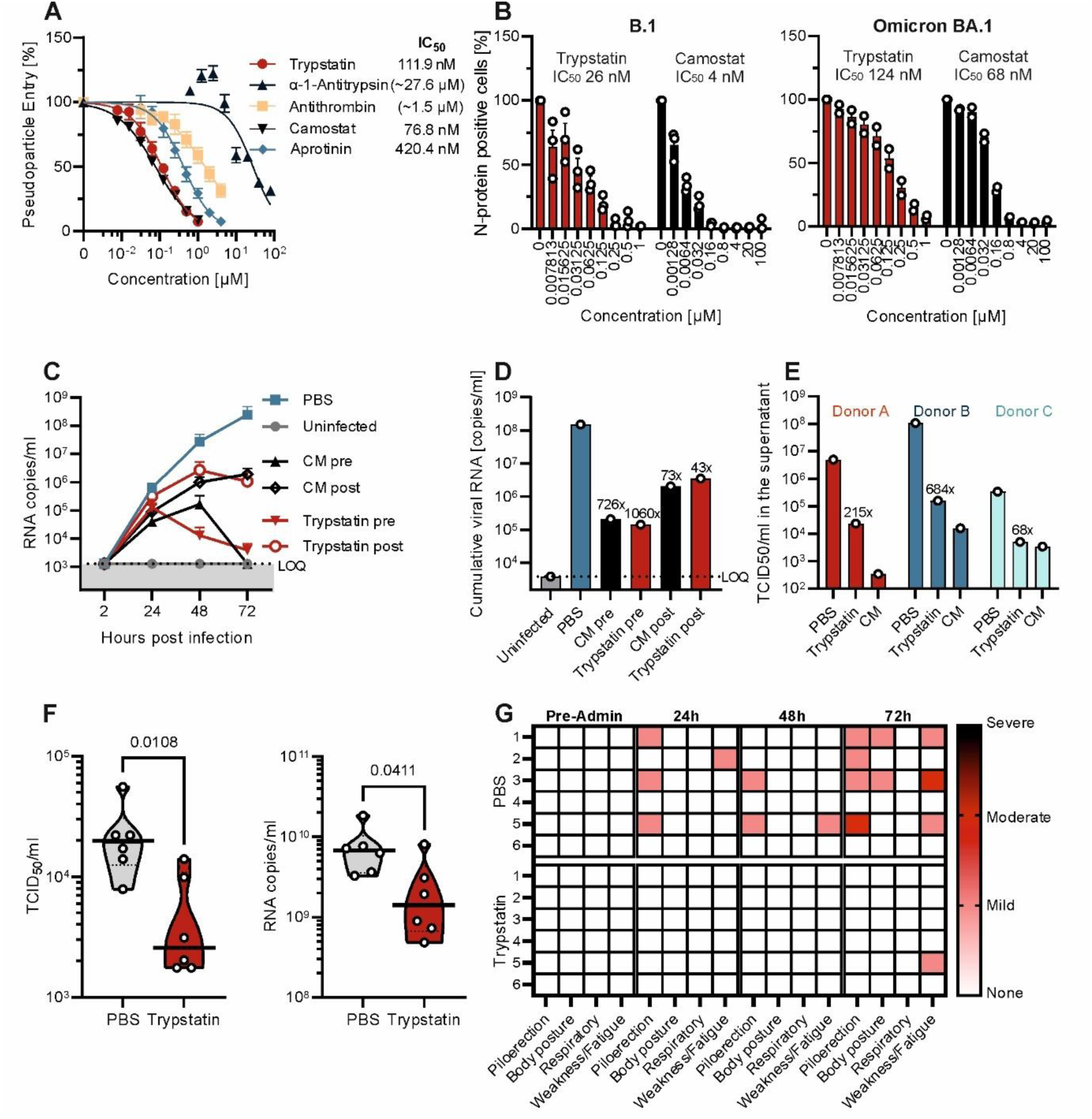
Trypstatin inhibits SARS-CoV-2 *in vitro* and *in vivo*. **(A)** Caco-2 cells were pre-treated with serial dilution of compounds before transduction with luciferase-encoding lentiviral pseudoparticles harboring the SARS-CoV-2 Hu-1 spike. Transduction rates were assessed 48 h later by measuring luciferase activity in cell lysates. Shown is the mean of three independent experiments performed in triplicates ± SEM. **(B)** Caco-2 cells were treated with indicated compounds for 30 min before inoculation with SARS-CoV-2 B.1 D614G (MOI 0.0005) or the Omicron BA.1 variant (MOI 0.2). Nucleocapsid protein positive cells were quantified one day post infection by flow cytometry. Shown are mean values of two (BA.1) or three (B.1) experiments performed in triplicates ± SEM. **(C)** Calu-3 cells were treated with Trypstatin (10 µM), camostat (80 μM) or PBS right before (pre) or 3 hours post (post) infection with SARS-CoV-2 BA.5 at a MOI of 0.001. Two hours later, the inoculum was removed, cells were washed and fresh medium supplemented with respective compounds was added. Complete supernatants were sampled at 2, 24, 48, and 72 h postinfection (hpi) and fresh medium with compound was added after sampling. RT-qPCR targeting SARS-CoV-2 ORF1b nsp14 was performed on RNA isolated from harvested supernatants. **(D)** Cumulative SARS-CoV-2 RNA copies until 3 dpi assessed by area under the curve analysis from **(C)**. **(E)** The apical site of human airway epithelial cells (HAEC) grown at air–liquid interface was exposed to PBS, Trypstatin (10 μM) or camostat mesylate (80 μM) before inoculation with SARS-CoV-2 Omicron BA. 5 (MOI 0.5) for 2 h before apical washing and further culturing at the air-liquid interface. Two days-post infection, mucus was washed off and PBS was added to the apical side for 30 minutes before the sample was subjected to TCID50 titration. Shown are TCID50/ml values of pooled apical washes from three cultures per donor. **(F)** Syrian gold hamsters were infected with SARS-CoV-2 (Delta) and treated with Trypstatin or PBS. They were sacrificed 72h post infection and bronchoalveolar lavage was collected for each animal. BAL samples were subjected to RT-qPCR and TCID50 analysis. Mann-Whitney-U-Test was applied to test for significance. **(G)** Hamsters were monitored daily for clinical symptoms.

To assess the stability and activity of Trypstatin in the presence of human airway mucus, we incubated Trypstatin in varying concentrations of HAEC-derived mucus for periods ranging from 0.5 to 4 hours before testing it on cells infected with SARS-CoV-2 pseudoparticles. Our results showed that Trypstatin fully retained its inhibitory capacity against spike-mediated entry, supporting its stability and sustained efficacy in mucus (Supplemental Figure 6).

Finally, we conducted an *in vivo* proof-of-concept study in a Syrian gold hamster model of SARS-CoV-2 infection. Control studies confirmed that Trypstatin effectively inhibits hamster TMPRSS2 with IC_50_ values similar to those for human TMPRSS2 (Supplemental Figure 7). For the *in vivo* study, six hamsters per group received a relatively low dose of 0.3 mg/kg of Trypstatin (or PBS as control) intranasally, administered 3 hours before and after inoculation with the highly pathogenic SARS-CoV-2 Delta variant, followed by daily dosing for two additional days (Supplemental Figure 8). Clinical symptoms were monitored daily and after 72 hours, animals were sacrificed, and bronchoalveolar lavage (BAL) fluid was collected for analysis. TCID_50_ and qPCR analyses of BAL samples revealed significantly reduced viral loads and infectious virus particles in the Trypstatin-treated group compared to the PBS-treated group (Figure 3F). Clinical observations showed that symptoms, such as piloerection, weakness, fatigue, and abnormal posture, were almost entirely absent in the Trypstatin-treated animals, in contrast to prominent symptoms in the PBS-treated group (Figure 3G). These findings confirm the *in vivo* efficacy of Trypstatin in reducing SARS-CoV-2 infection and associated clinical symptoms in a relevant animal model.

### Broad-spectrum inhibition of corona- and influenzaviruses by Trypstatin

Besides SARS-CoV-2, other human coronaviruses also rely on TMPRSS2 or its paralogs for host cell entry (19, 58, 59). To assess Trypstatin’s effectiveness against these viruses, we conducted *in vitro* assays using pseudoparticles carrying spikes from SARS-CoV-1 and MERS-CoV, as well as replication-competent hCoV-NL63. Trypstatin demonstrated dose-dependent inhibition, with IC_50_ values of 3 nM for SARS-CoV-1 and 78 nM for MERS-CoV (Figure 4A-B). In addition, Trypstatin effectively reduced replication of hCoV-NL63, with an IC_50_ of 195 nM (Figure 4C), highlighting its broad-spectrum activity across both alpha- and beta-coronaviruses. Since TMPRSS2 and related proteases are also utilized by influenza viruses to activate hemagglutinin for host cell fusion (17, 22, 23), we evaluated Trypstatin’s activity against influenza A and B viruses. Human lung cells (Calu-3) were pre-treated with varying concentrations of Trypstatin, followed by inoculation with two predominant circulating influenza A subtypes (H1N1 and H3N2) and one influenza B virus strain. Trypstatin inhibited all three viruses in a dose-dependent manner, with IC_50_ values of approximately 292 nM for H1N1, 144 nM for H3N2, and 570 nM for influenza B (Figure 4D-F). In summary, Trypstatin demonstrates broad inhibition of coronaviruses and currently predominant influenza viruses in human lung cells.

**Figure 4:**
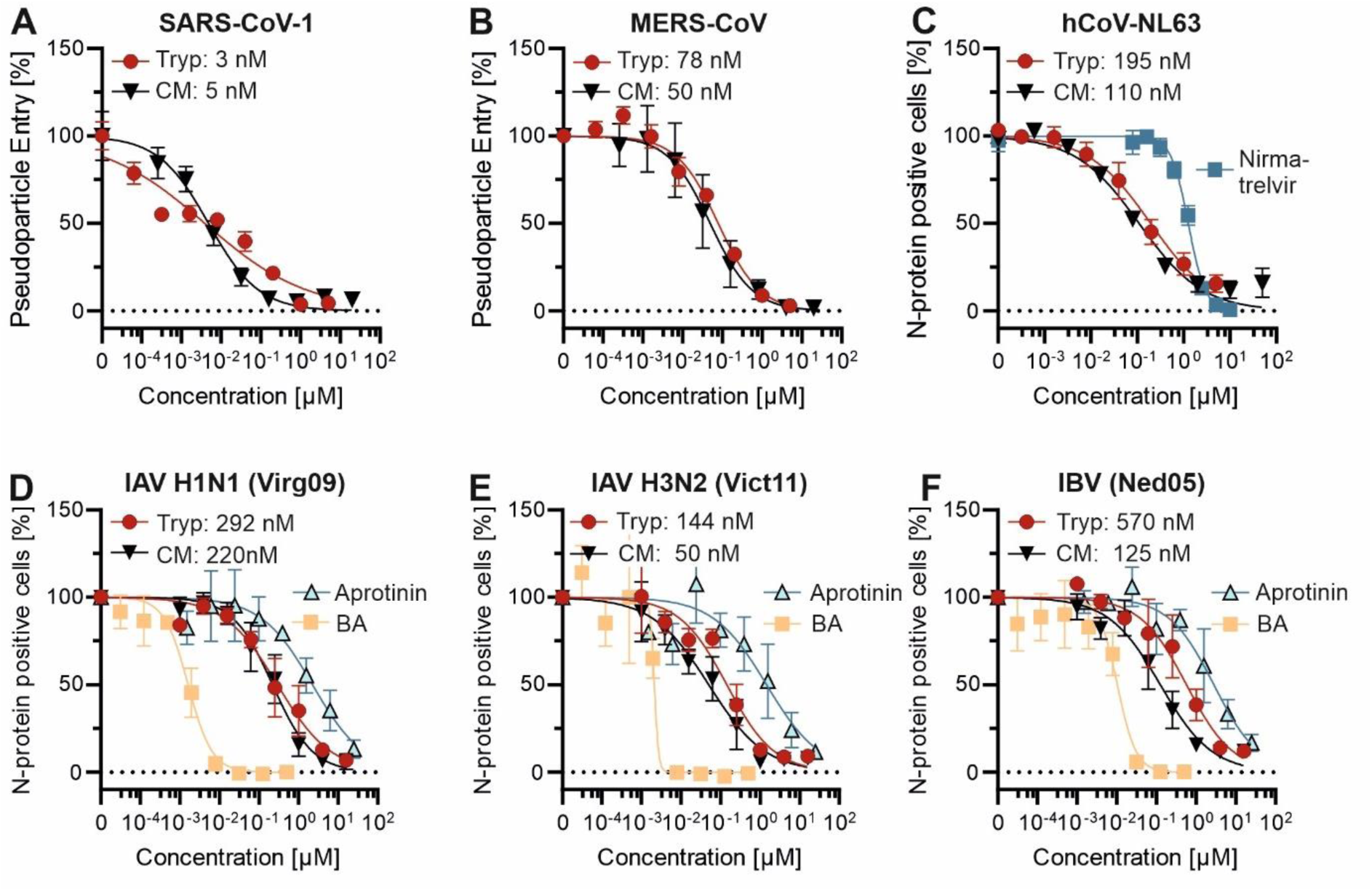
Trypstatin is a broad-spectrum inhibitor of corona- and influenza viruses. **(A-B)** Caco-2 cells were treated with serial dilutions of compound right before transduction with lentiviral pseudoparticles harboring the SARS-CoV-1- or MERS-CoV spike. Transduction rates were assessed 48 h later by measuring luciferase activity in cell lysates. **(C)** Caco-2 cells were treated with serial dilutions of Trypstatin or control inhibitors before infection with hCoV-NL63 (MOI 0.018) virus. Nucleocapsid protein positive cells were quantified three days post infection by flow cytometry. **(D-F)** Calu-3 cells were exposed to serial dilution of compounds for 30 minutes before inoculation with 100xTCID50 per well of respective influenza virus. Three days post-infection, nucleocapsid positive cells were quantified by immunostaining and high-content imaging. Shown are mean values of three independent experiments performed in triplicates ± SEM for all graphs. Tryp = Trypstatin, CM = Camostat mesylate, BA = Baloxavir acid.

## 4. Discussion

Our findings identify and characterize Trypstatin as a promising broad-spectrum antiviral candidate targeting TMPRSS2-dependent viruses, including SARS-CoV-2 and influenza virus. The efficacy of Trypstatin across various virus species, its stability in mucus-rich environments, and favorable activity in an animal model underscore its potential as an effective antiviral agent against respiratory infections.

Bikunin, the precursor of the Trypstatin fragment, is a multifunctional glycosylated protease inhibitor and a key member of the inter-α-inhibitor family. It is present in plasma at baseline concentrations of approximately 2.5 µg/mL (0.1 µM), which can increase to 36 µg/mL (1.4 µM) under inflammatory conditions (60). While Bikunin regulates inflammation, immune responses, and specifically protease activity through its two Kunitz-type domains, its antiviral potential in the full-length form is constrained by steric hindrance. The first Kunitz domain (residues 231-281) partially obstructs the P1-containing loop of the second domain (residues 287-337), reducing its ability to interact with larger substates than trypsin (39, 61). In contrast, Trypstatin (residues 284-344) and the second Kunitz domain exhibit potent antiviral activity against TMPRSS2 when functioning as independent polypeptides. To date, Trypstatin has only been described as a distinct protein in rats (50, 51), and its presence and activity in human plasma remains speculative. Theoretically, full proteolytic processing of Bikunin could produce sufficient plasma concentrations of Trypstatin to exert antiviral effects. This cleavage is thought to occur under conditions such as inflammation, tissue injury, or infections, where proteolytic enzymes like trypsin, elastase, or plasmin are elevated (60). Moreover, localized proteolytic activity in respiratory tissues or epithelial barriers may further enhance the activation of Trypstatin in response to viral challenges. However, the physiological significance of this activation remains to be determined.

TMPRSS2, a transmembrane serine protease, plays a vital role in the entry of respiratory viruses such as SARS-CoV-2 and influenza by cleaving and activating viral glycoproteins essential for membrane fusion (20, 21, 23, 52). We show that Trypstatin effectively blocks TMPRSS2 by binding to the S1 specificity pocket of the protease, thus inhibiting enzymatic activity and thus the cleavage step, thereby preventing viral entry. Molecular dynamics simulations provide insights into the structural affinity of Trypstatin for TMPRSS2, revealing strong electrostatic interactions within the S1 pocket that stabilize the binding of the inhibitor. The specific configuration of the P1-positioned arginine residue in Trypstatin competes effectively with viral glycoproteins, and likely block access to the active site of the protease. This structural specificity suggests that Trypstatin can serve as a targeted TMPRSS2 inhibitor, offering comparable efficacy to small-molecule inhibitors such as camostat mesylate while benefiting from the lower cytotoxicity and tolerability often associated with endogenous peptides (62).

TMPRSS2 is a highly promising target for antiviral therapy due to several distinctive characteristics. First and foremost, it plays a critical role in viral entry for various virus families, including coronaviruses, influenza viruses, and certain paramyxoviruses (63, 64). Notably, Trypstatin also inhibits TMPRSS11D and TMPRSS13, two proteases involved in the cleavage and activation of viral glycoproteins. This suggests that its inhibitory activity extends to related proteases essential for the entry of other respiratory viruses (26, 27). For these viruses, TMPRSS2 and related proteases cleave envelope glycoproteins - such as the spike protein in coronaviruses and hemagglutinin in influenza - triggering the conformational changes necessary for membrane fusion with host cells (20–22, 52, 63). Thus, inhibiting TMPRSS2 is an effective strategy for blocking viral infection at its earliest stage (18). A key advantage of targeting TMPRSS2, a host protein, is its resilience against viral mutations that often drive resistance to conventional antivirals (65). Since viruses depend on TMPRSS2 for entry, they are less likely to evade its inhibition through mutations. This host-targeting approach provides a durable defense against viral mutations and emerging variants, as demonstrated by Trypstatin’s potent inhibition of all analyzed SARS-CoV-2 variants of concern (VOCs). Moreover, studies in TMPRSS2 knockout mice have shown reduced susceptibility and pathology following infection with influenza viruses or SARS-CoV-2, including highly transmissible variants such as Omicron (66–68).

The expression of TMPRSS2, TMPRSS11D, and TMPRSS13 in human respiratory tract epithelial cells - particularly in lung tissues where respiratory viruses establish infection - further underscores its importance as a drug target (69). TMPRSS2 inhibition could provide localized protection, focusing directly on the tissues most affected by respiratory viruses. Importantly, inhibition of TMPRSS2 has minimal impact on essential cellular processes. TMPRSS2 knockout mice develop without significant clinical phenotypes and remain fertile, highlighting the selective effect of TMPRSS2 inhibition on viral entry while sparing critical cellular functions (70).

Interestingly, anti-proteolytic activity of Trypstatin also extends to neutrophil elastase and kallikrein 5. Neutrophil elastase contributes to acute respiratory distress and amplifies pro-inflammatory cascades (54, 55), while kallikrein exhibits similar functions and was recently identified as a priming protease for the spike protein of various beta-coronaviruses (56, 71). This raises the possibility that Trypstatin may exhibit mild anti-inflammatory properties, although its effects on these targets are less pronounced due to significantly higher IC_50_ values compared to its inhibition of transmembrane serine proteases.

The broad-spectrum antiviral activity of Trypstatin against SARS-CoV-2, SARS-CoV-1, MERS-CoV, hCoV-NL63, and influenza A and B underscores its potential as a versatile treatment for respiratory pathogens. This makes it particularly valuable for addressing seasonal influenza strains, where vaccination is often less effective due to different circulating variants, as well as for emerging viruses that demand rapid therapeutic solutions. *In vitro* studies show that Trypstatin reduces viral replication in human lung epithelial cells at nanomolar concentrations, demonstrating its potency as an inhibitor. Importantly, Trypstatin remains active in mucus - a common barrier that can diminish drug efficacy (72), making it well-suited for real-world respiratory applications. *In vivo* studies in the Syrian gold hamster model further highlight its efficacy, with low-dose treatment providing significant antiviral effects without observed side effects. This is in stark contrast to other TMPRSS2 inhibitors, which required intranasal administration at concentrations up to 100 times higher under similar conditions (73–75). While these results are promising, further studies are necessary to explore the effects of higher Trypstatin concentrations to fully assess its therapeutic range and safety profile.

Kunitz-type inhibitors have demonstrated clinical viability, with examples such as ecallantide, approved for hereditary angioedema treatment, and others such as Depelestat, Amblyomin-X, and Sivelestat advancing to clinical trials (55, 76–78). These successful examples highlight their scalability and therapeutic potential. Notably, aprotinin, another Kunitz-type inhibitor, reached phase III trials as a COVID-19 therapy and demonstrated the ability to inhibit TMPRSS2 and exhibit antiviral activity *in vitro* (79–82). However, aprotinin, derived from bovine lung tissue, was temporarily withdrawn from the European market due to severe side effects, including immunogenicity, allergic reactions, and off-target effects associated with its broad-spectrum antiprotease activity (83). These limitations emphasize the need for safer and more targeted alternatives. Trypstatin offers distinct advantages over aprotinin. As an endogenous human protein fragment derived from Bikunin, Trypstatin is expected to have significantly lower immunogenicity and a superior safety profile. Its high specificity for TMPRSS2 and related proteases, such as TMPRSS11D and TMPRSS13, minimizes off-target effects, while its stability in mucus in the respiratory tract makes it particularly well-suited for intranasal or inhalable applications. The high specificity of Trypstatin and potent inhibition of TMPRSS2 further distinguish it from other protease inhibitors, such as antithrombin and α1-antitrypsin, which exhibit lower specificity and higher IC_50_ values (30, 49). This exceptional potency, coupled with its host-targeting mechanism, reduces the risk of resistance and minimizes drug-drug interactions. Unlike existing antiviral therapies such as Paxlovid, which face contraindication challenges (84), Trypstatin provides a safer and more adaptable solution. Its combination of specificity, efficacy, scalability, and safety positions Trypstatin as a promising candidate for respiratory viral infections in both prophylactic and therapeutic applications.

## Data availability statement

The data that support the findings of this study are available from the corresponding author (J.M.) upon request.

## Ethic statements

Ethic approval for the generation of peptide libraries from hemofiltrate was obtained from the Ethics Committee of Ulm University (91/17). Collection of tissue samples for the generation of human airway epithelia cell cultures has been approved by the ethics committee at the Medical School Hannover (airway tissue, application number 2699-2015). The housing of animals was in accordance with EU Directive 2010/63/EU of 22 September 2010 on the protection of animals used for scientific purposes. All animal experimentation was carried out under a license approved by the Swedish Board of Agriculture and the regional animal ethics committee in Stockholm. The study was performed under the animal ethical permit number: 2020-2021.

## Acknowledgments

We thank Daniela Krnavek, Nicola Schrott, Carolin Weiss, Tanja Tubbesing and Isabell Haußmann for their technical assistance.

## Supporting Information

**Supplemental Figure 1:**
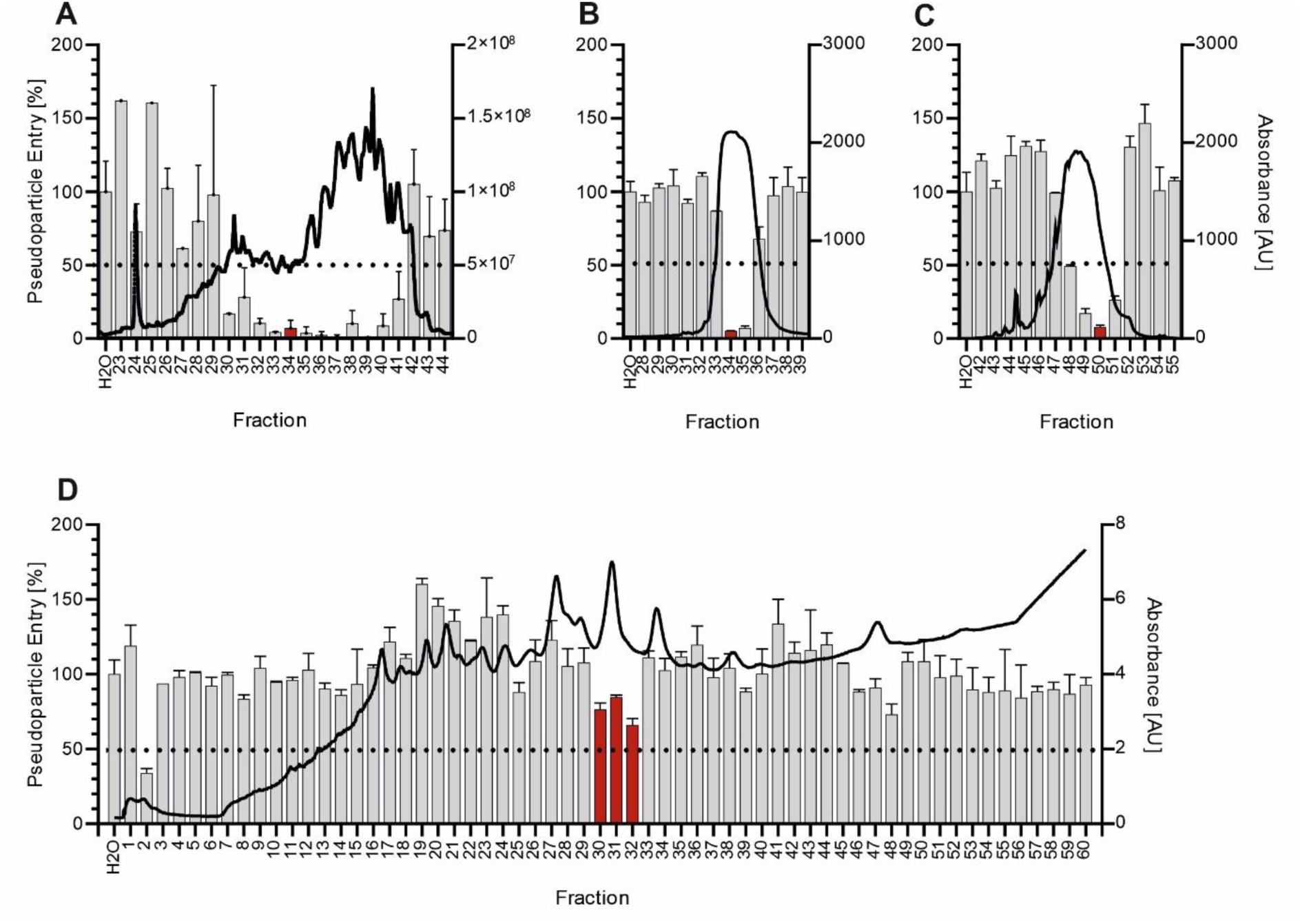
High-throughput screening for endogenous inhibitors of SARS-CoV-2 entry. **(A-D)** Caco-2 cells were pre-treated with peptide/protein containing fractions of subpurified hemofiltrate library fractions before transduction with luciferase-encoding lentiviral pseudoparticles harboring the SARS-CoV-2 Hu-1 spike. Transduction rates were assessed 48 h later by measuring luciferase activity in cell lysates. Columns represent pseudoparticle entry and black line absorbance at 280 nm of the corresponding fraction. Fractions marked in red (A-C) were subpurified before final fractions **(D)** were analyzed by LCMS/MS. Shown are mean values of one experiment performed in triplicates ± SEM.

**Supplemental Figure 2:**
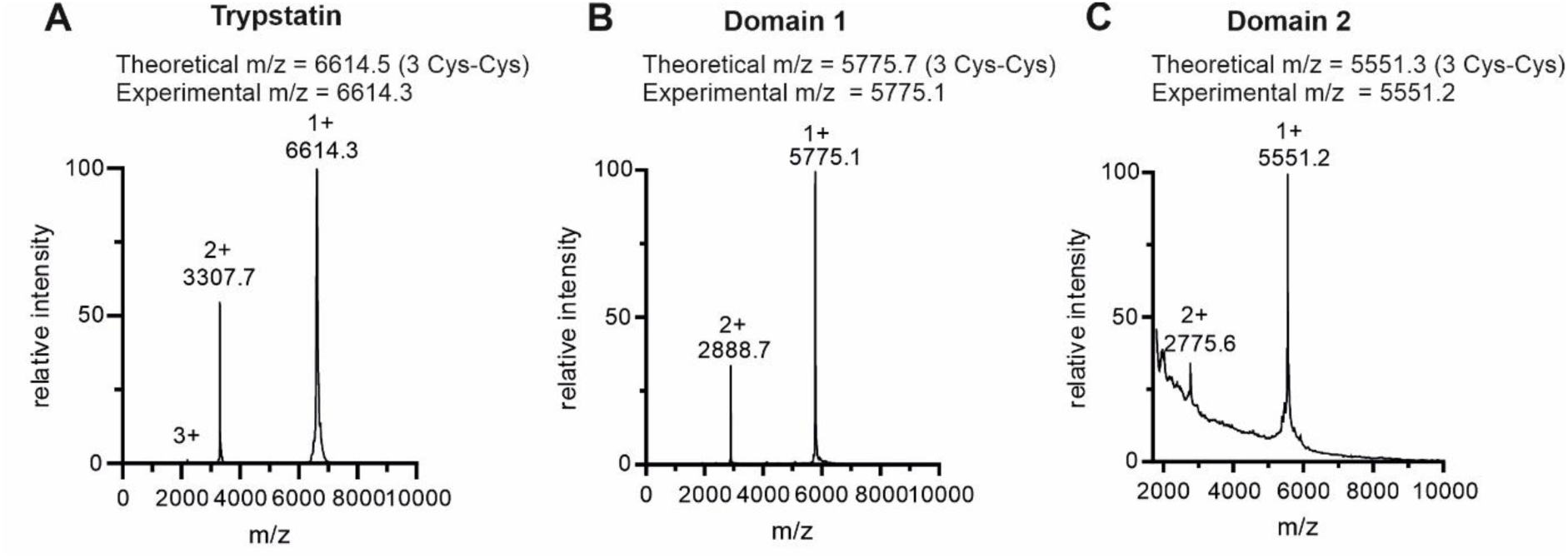
MALDI-TOF-spectra of synthetically produced and refolded kunitz domains. **(A)** Trypstatin (Protein AMBP, P02760: residues 284-344), **(B)** BPTI/Kunitz inhibitor 1 domain (Protein AMBP, P02760: residues 231-281), (C) BPTI/Kunitz inhibitor 2 domain (Protein AMBP, P02760: residues 287-337). In all cases, the experimental average masses indicated the formation of three disulfide bridges.

**Supplemental Figure 3:**
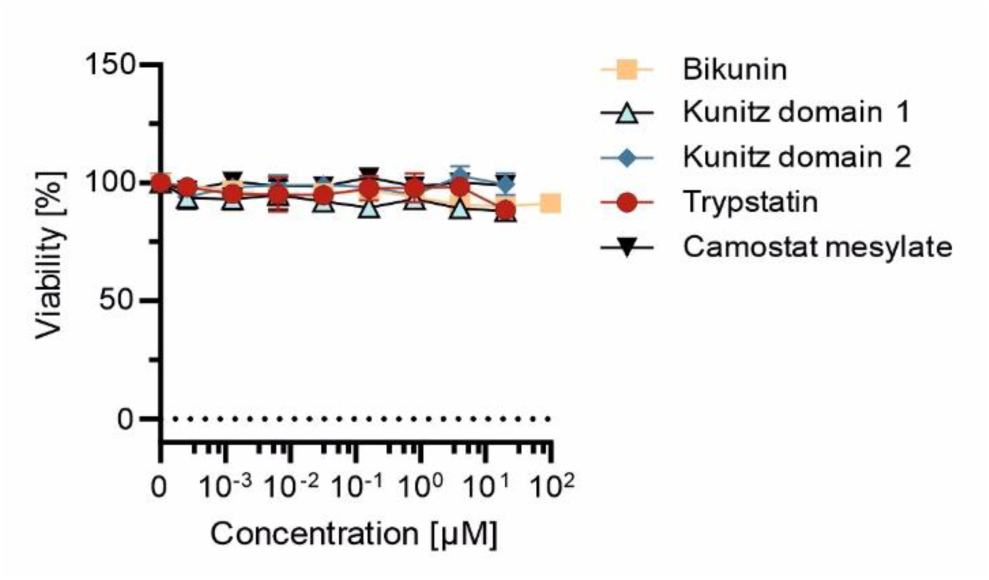
Trypstatin does not influence cell viability. Caco-2 cells were treated with serial dilutions of compounds for 48 h before cell viability was assessed using the Promega CellTiter-Glo® Cell Viability Assay. Shown are mean values of three independent experiments performed in triplicates ± SEM.

**Supplemental Figure 4:**
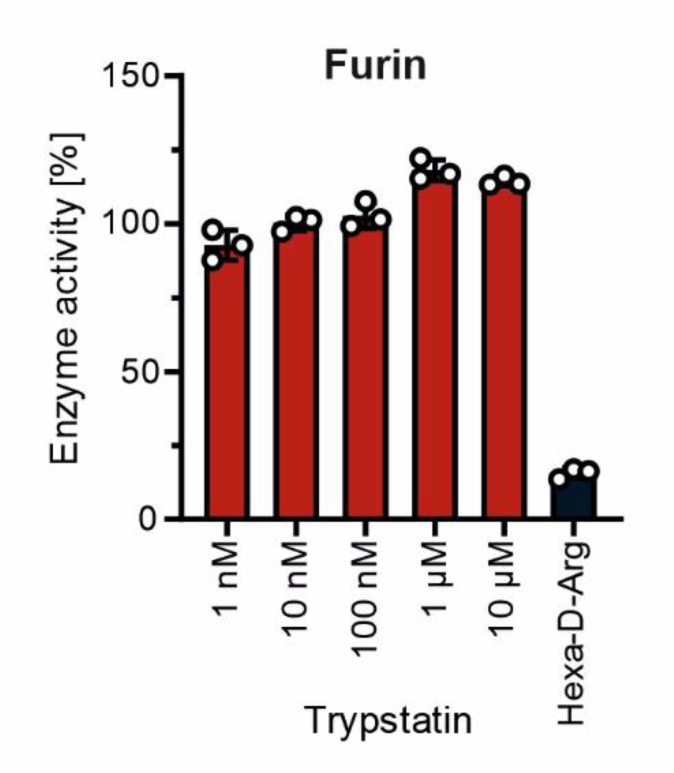
Trypstatin does not inhibit furin. Compounds were mixed with recombinant furin before addition of a fluorogenic reporter substrate. Fluorescence intensity was measured at an excitation wavelength of 355 nm and emission wavelength of 460 nm. Shown are mean values of three independent experiments performed in duplicates ± SEM.

**Supplemental Figure 5:**
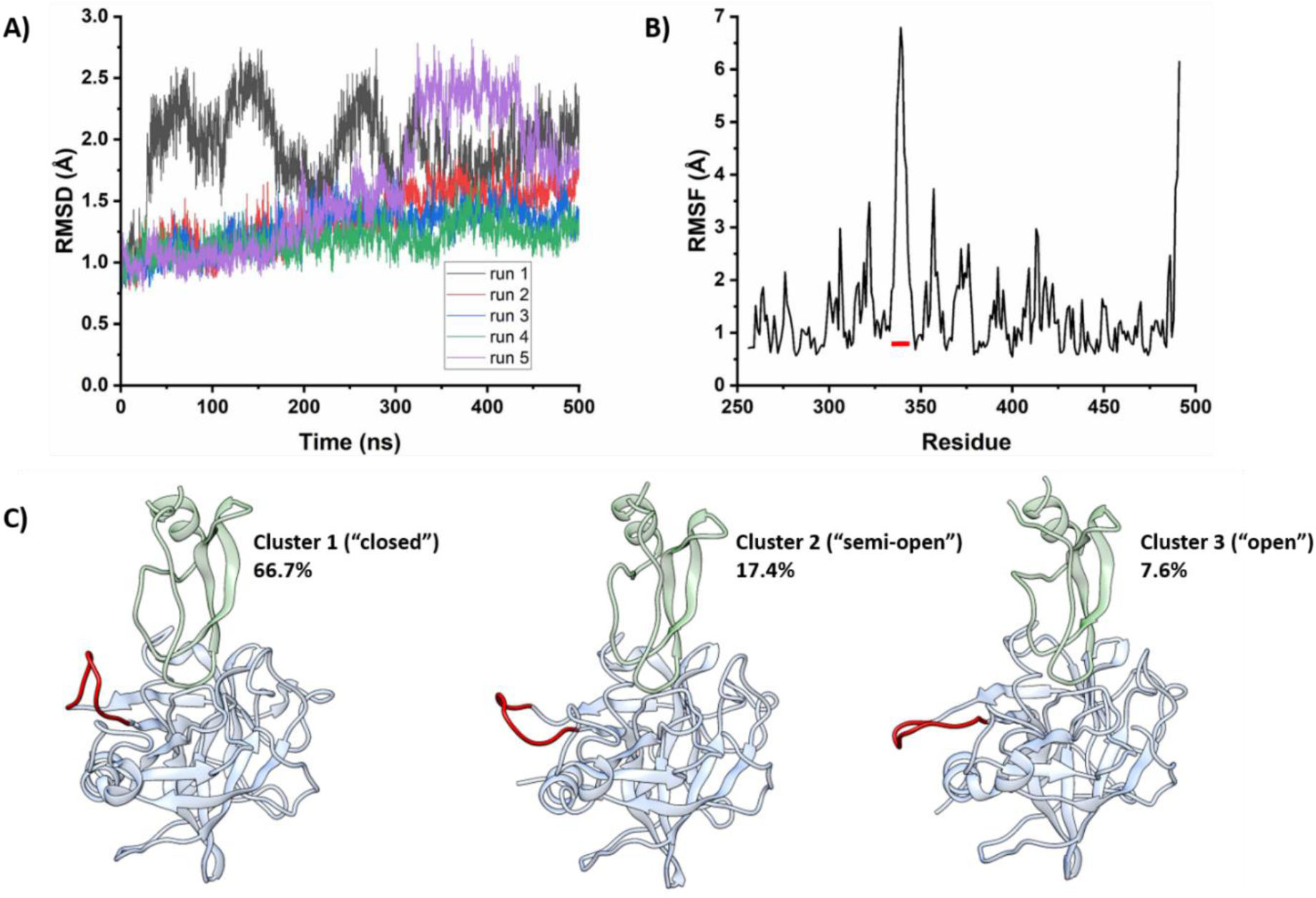
The simulations showed the dynamic behavoiur of the backbone atoms of TMPRSS2. **(A)** RMSD evolution of the backbone atoms of TMPRSS2. (**B)** The RMSF analysis shows the high flexibility of the loop (highlighted by the red line) encompassing residues 336-345 of TMPRSS2. (**C)** Representative structures from the clustering of the TMPRSS2’s 336-345 loop (red). The clustering analysis of this loop showed three clusters of structures, which can be classified according to the relative orientation with respect to Trypstatin. The percent refers to the cluster size of the sampled population during 1.5 µs. The most populated cluster of structures (66.7%, the loop is highlighted in red), displays a “closed” conformation of the loop (“near” Trypstatin), followed by a cluster with the loop positioned on an intermediate orientation with respect to Trypstatin (semi-open, 17.4%), and other cluster of structures with the loop adopting an “open” conformation, further away from Trypstatin (7.6%). Although “open” complexes represented only 7.6% of the total frames, they were prevalent in replicas 1 and 5 of the simulations. RMSD = root mean sqare deviation, RMSF = root mean square fluctuation.

**Supplemental Figure 6:**
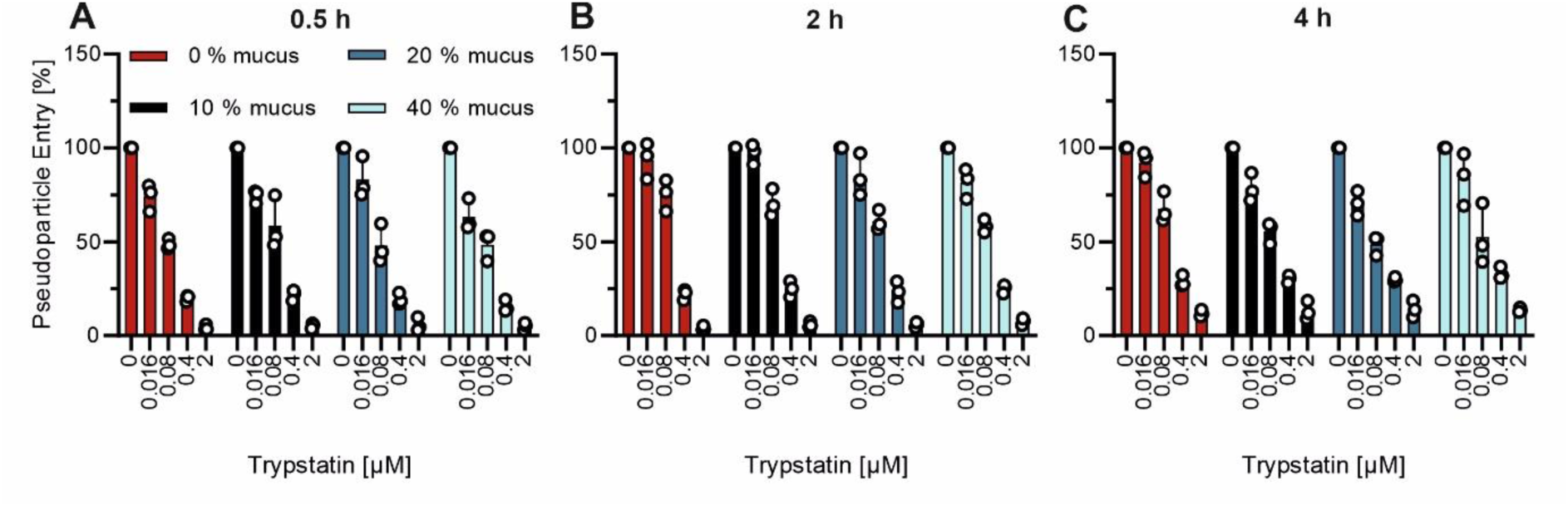
Trypstatin remains active in the presence of human airway mucus. Serial dilutions of Trypstatin were mixed to the indicated concentrations with mucus from HAEC cultures at 37°C for 30 min (A), 2 h (B) or 4 h (C) before addition to Caco-2 cells and subsequent transduction with luciferase encoding lentiviral pseudoparticles harboring the SARS-CoV-2 Hu-1 spike. Transduction rates were assessed 48 h later by measuring luciferase activity in cell lysates. Shown are mean values of experiment performed in triplicates ± SEM.

**Supplemental Figure 7:**
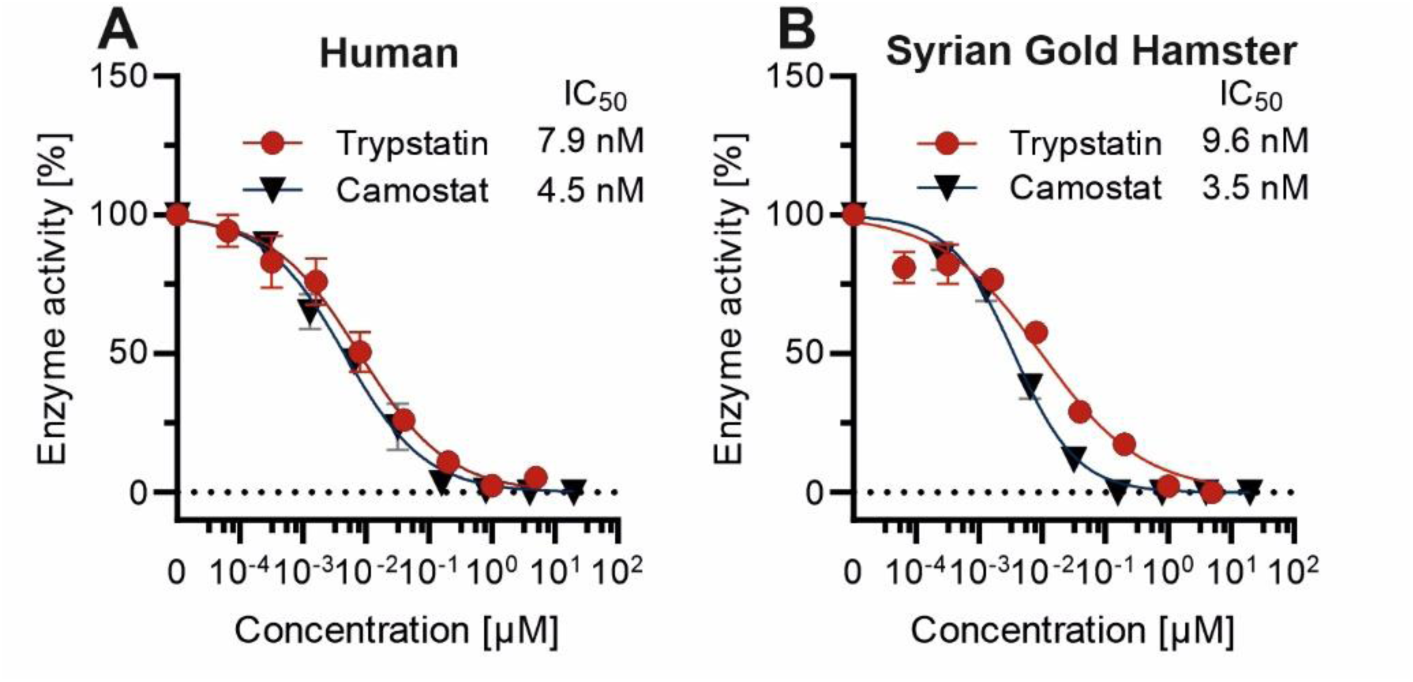
Trypstatin blocks hamster and human TMPRSS2 activity with similar efficacies. HEK293T-cells were transfected with human or hamster TMPRSS2 expression plasmid or mock control, respectively, before treatment with serial dilutions of inhibitors and the addition of a fluorogenic reporter substrate. Values were corrected for the signal of mock-transfected HEK293T-cells. Shown are mean values of one experiment performed in triplicates ± SEM.

**Supplemental Figure 8:**
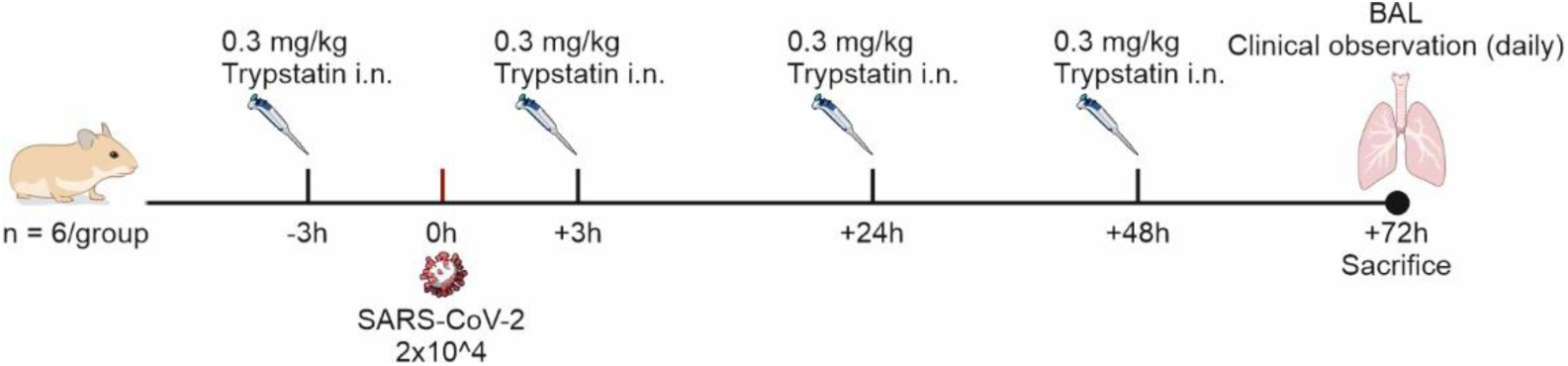
Treatment schedule of *in vivo* study. 12 syrian gold hamsters were randomized into groups of 6. Groups were either treated with 0.3 mg/ml Trypstatin or PBS at the indicated time points. Hamsters were inoculated with the SARS-CoV-2 delta variant. 72h post-infection, hamsters were sacrified and bronchoalveolar lavage was taken. Clinical symptoms were recorded on a daily basis.

